# The Mitochondrial Brown Adipose Tissue Maintenance Factor Nipsnap1 Interfaces Directly with the Beta-Oxidation Protein Machinery

**DOI:** 10.1101/2024.12.29.630678

**Authors:** Pei-Yin Tsai, Yue Qu, Claire Walter, Yang Liu, Chloe Cheng, Joeva J Barrow

## Abstract

**Background:** The activation of brown adipose tissue (BAT) is associated with improved metabolic health in humans. We previously identified the mitochondrial protein 4-Nitrophenylphosphatase Domain and Non-Neuronal SNAP25-Like 1 (Nipsnap1) as a novel regulatory factor that integrates with lipid metabolism and is critical to sustain the long-term activation of BAT, but the precise mechanism and function of Nipsnap1 is unknown.

**Objectives:** Define how the regulatory factor Nipsnap1 integrates with lipid metabolism.

**Methods:** We generated adeno-associated viral (AAV) constructs that overexpress Nipsnap1 in the thermogenic fat of mice. We then measured both whole-body and cellular mitochondrial metabolism and mapped the first Nipsnap1 interacting protein-protein network.

**Results:** Herein, we show that adipose-specific overexpression of Nipsnap1 in mice increases energy expenditure through the utilization of lipids as an energy substrate. The increase in energy expenditure results in reduced weight gain. Additionally, we show that Nipsnap1 overexpression in primary adipocytes increases lipid beta-oxidation. Moreover, we mapped the first protein-protein network of Nipsnap1 in brown adipocytes and show that Nipsnap1 interacts with proteins that regulate both peroxisomal and mitochondrial fatty acid beta-oxidation.

**Conclusion:** This study elucidates a mechanistic function of Nipsnap1 in thermogenic fat where Nipsnap1 facilitates a functional connection between peroxisomal and mitochondrial beta-oxidation pathways. By enhancing lipid utilization as energy substrates, Nipsnap1 plays a pivotal role in sustaining thermogenic fat activation to increase energy expenditure. These findings underscore the potential of Nipsnap1 as a therapeutic target for metabolic health.

## INTRODUCTION

The activation of human brown adipose tissue (BAT) to enhance cardiometabolic health remain an attractive molecular mechanism to combat obesity-related metabolic dysfunction^1–6^. One of the challenges surrounding brown fat activation however is that activation of this protective metabolic depot is extremely transient in nature and the regulatory factors required to sustain the activation of the BAT depot is poorly understood^7^. We have previously defined a novel BAT maintenance regulatory factor termed 4-Nitrophenylphosphatase Domain and Non-Neuronal SNAP25-Like 1 (Nipsnap1) that is critical for the sustained activation of BAT^8^. We demonstrated that Nipsnap1 localizes to the mitochondrial matrix in brown fat and increases its transcript and protein levels in response to both chronic cold and pharmacological β3 adrenergic signaling. Through the generation of the first BAT-specific Nipsnap1 KO mice, we demonstrated that although the metabolic activation of BAT to increase energy expenditure was intact, the Nipsnap1 knockout mice were unable to sustain this activation in the face of an extended environmental cold challenge. Mechanistically, we demonstrated that ablation of Nipsnap1 drove the inability to sustain BAT activation which was caused by severe defects in lipid metabolism and beta-oxidation capacity. How Nipsnap1 integrates with lipid metabolism however, and the molecular protein binding targets and network for Nipsnap1 in brown fat is currently unknown.

Herein, we map the first Nipsnap1 protein network in brown fat and show that Nipsnap1 directly binds to proteins involved in mitochondrial and peroxisomal beta-oxidation. We further demonstrate that overexpression of Nipsnap1in brown fat can enhance beta-oxidation capacity and increase energy expenditure in mice leading to protective metabolic health benefits. Taken together, Nipsnap1 plays a critical role in the maintenance of BAT and offers a new avenue for therapeutic opportunities to enhance metabolic health.

## RESULTS

### Overexpression of Nipsnap1 enhances protection against environmental cold challenge

Ablation of Nipsnap1 causes defects in the sustained activation of brown fat leading to lowered energy expenditure and impaired lipid metabolism which results in failures to protect body temperature in the face of a cold- environmental challenge^8^. To investigate if elevated Nipsnap1 protein levels in brown adipocytes would confer enhanced metabolic benefits, we engineered adipose-specific Nipsnap1 adeno-associated virus serotype 8 (AAV8) constructs. These constructs included a Flag-tagged Nipsnap1 overexpression vector and a GFP control vector (Figure 1A). These constructs were tail vein injected into C57BL6/J wildtype mice and overexpression was confirmed by gene and protein expression in BAT tissue 3 weeks post infection (Figure 1B and 1C). Overexpression (OE) of Nipsnap1 was confirmed to be predominantly expressed BAT and iWAT adipose tissue depots as there was no expression of Flag tagged Nipsnap1 in the liver or brain where Nipsnap1 is predominately expressed (Figure 1D and 1E). Mitochondrial localization of our Flag-tagged Nipsnap1construct was also confirmed by BAT fractionation (Figure 1F). Functionally, we discovered that Nipsnap1 OE mice had higher core body temperature when faced with a cold challenge and gained less weight during the exposure period compared to control mice (Figure 1G and 1H). This protection occurred with no changes in UCP1 protein or RNA levels which is consistent with our previous findings with Nipsnap1 ablation, where the significant defects in body temperature, energy expenditure, and lipid metabolism were independent of changes with UCP1 protein and RNA levels (Figure 1C and 1I)^8^.

**Figure 1.**
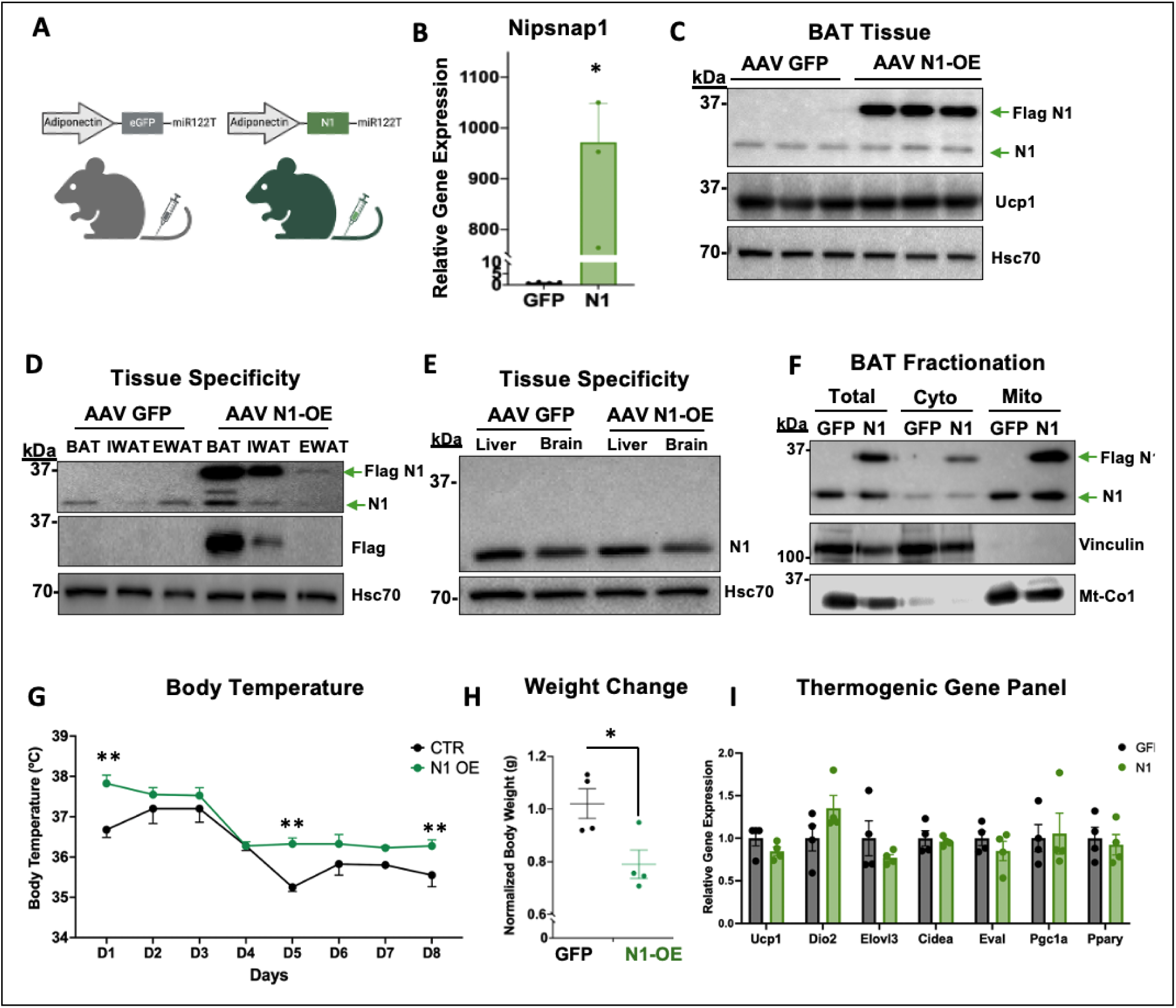
Overexpression of Nipsnap1 enhances protection against environmental cold challenge. **(A)** Schematic of the AAV Nipsnap1 (N1) overexpression plasmid. **(B)** Relative gene expression of N1 in brown adipose tissues of male mice transduced with AAV-GFP or AAV-Flag-B1 at a multiplicity of infection (MOI) of 2.5x10^12 during a chronic cold exposure (n=3). **(C)** Representative Western blot illustrating the expression of endogenous N1 and AAV-Flag-N1 overexpression in brown adipose tissues (n=3). **(D, E)** Representative Western blots illustrating the specificity of AAV-Flag-N1 expression in brown (BAT), inguinal (IWAT), and white (EWAT) adipocytes, with no expression in the liver and brain. **(F)** Fractionation analysis of endogenous N1 and exogenous AAV-Flag-N1 in brown adipose tissue. Vinculin- loading control for cytoplasmic proteins, Mt-Co- loading control for mitochondrial proteins. **(G)** Daily rectal temperatures of male mice transduced with AAV-GFP or AAV-Flag-N1 in chronic cold exposure (n=4). **(H)** Relative body weight changes of male mice transduced with AAV-GFP or AAV-Flag- N1 after 8 days of cold exposure (n=4). **(I)** Relative gene expression of various thermogenic genes in brown adipose tissues of male mice transduced with AAV-eGFP or AAV-Flag-N1 after 8 days of cold exposure (n=4). All figures and data are represented as mean ± SEM. Significance is expressed as * p < 0.05, ** p < 0.01, *** p < 0.001 by Student’s t-test.

### Overexpression of Nipsnap1 increases energy expenditure in mice by increasing lipid substrate utilization

The increase in body temperature prompted the investigation in whether there was also a correspondent increase in energy expenditure. We therefore placed Nipsnap1 OE and control mice under cold-exposed conditions in Promethion metabolic cages to measure whole-body metabolism. We first assessed respiratory exchange ratio (RER) to examine macronutrient fuel utilization and discovered that Nipsnap1 OE mice had significantly lower RER rates which is indicative of increased lipid utilization predominantly in the night cycle when the mice are most active (Figure 2A and B). This indicated that the overexpression of Nipsnap1 led to a metabolic shift to metabolizing lipids as their primary energy source over carbohydrates. These findings are in contrast to the effect that was seen with our BAT-specific Nipsnap1 knockout mice, where we demonstrated previously that ablation of Nipsnap1 led to significant increases in RER due to impaired lipid metabolism and utilization^8^. The increase in lipid utilization of the Nipsnap1 OE mice compared to controls corresponded to significant increases in whole body energy expenditure with no changes in food intake or locomotive movement (Figure 2C-H). Taken together, our data suggests that Nipsnap1 is essential in the regulation of lipid metabolism.

**Figure 2.**
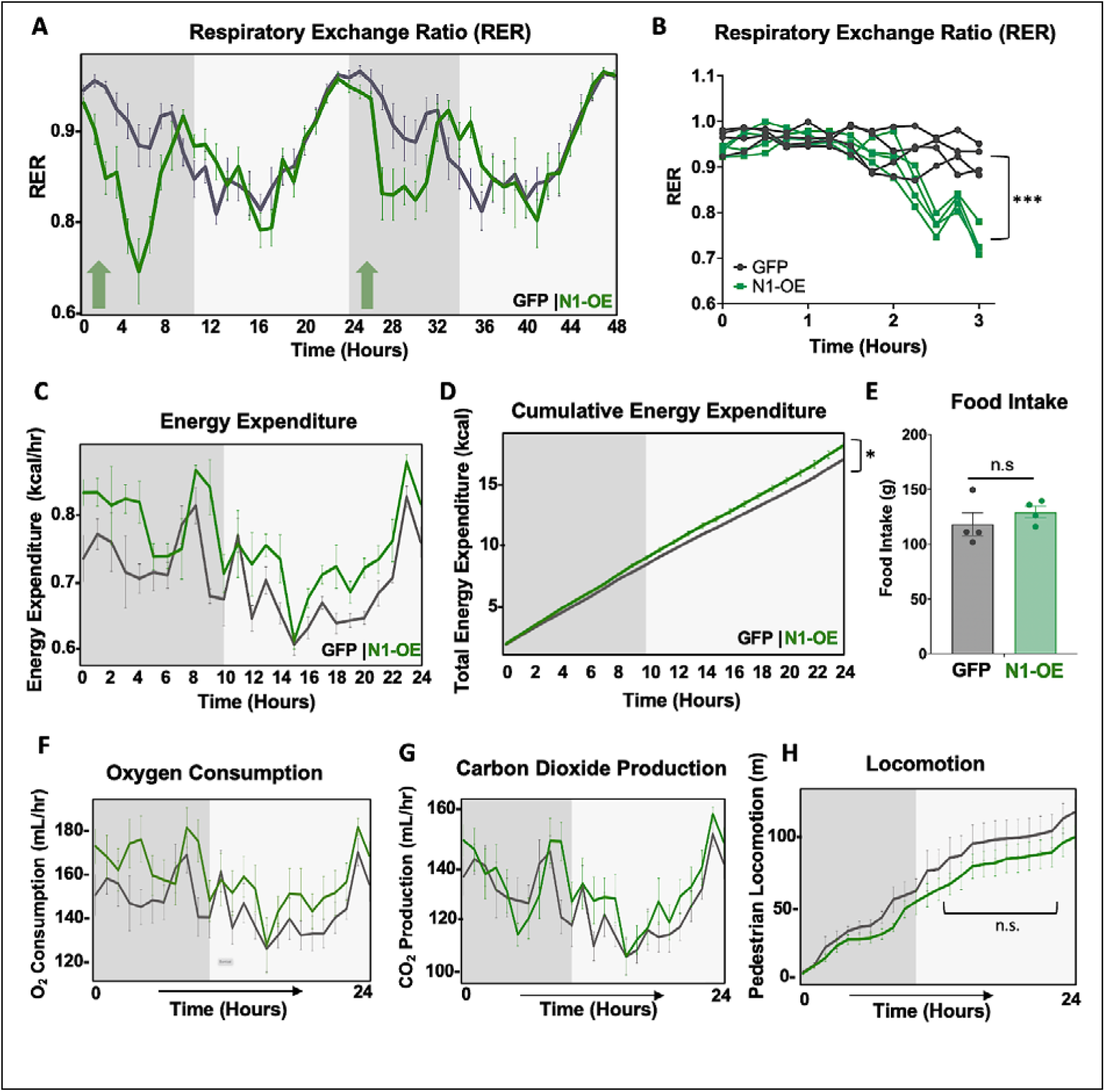
Overexpression of Nipsnap1 increases energy expenditure in mice by increasing lipid substrate utilization. Male mice transduced with AAV-GFP or AAV-Flag-N1 after 8 days of chronic cold exposure. Grey areas represent the dark cycle, and white areas represent the light cycle (n = 4). **(A, B)** Respiratory exchange ratio, **(C)** Energy expenditure, **(D)** Cumulative energy expenditure, **(E)** Total food intake, **(F)** Oxygen consumption, **(G)** Carbon dioxide production, **(H)** and locomotor activity. Whole body metabolic measurements were analyzed by ANCOVA. All other figures unless otherwise indicated are data represented as mean ± SEM. ∗p < 0.05, ∗∗p < 0.01 ∗∗∗p < 0.001 by Student’s t test.

### Overexpression of Nipsnap1 increases mitochondrial beta-oxidation capacity in brown adipocytes

To determine if the increase in lipid utilization in mice is driven primarily by brown adipocytes, we treated primary brown adipocytes with AAV-GFP or AAV-Nipsnap1 OE constructs (Figure 3A). We show that cells that overexpress Nipsnap1 had significantly higher rates of lipid beta-oxidation capacity compared to GFP treated controls (Figure 3B and C). We also observed trending increases in mitochondrial oxygen consumption when Nipsnap1 OE cells were stimulated with the β3 agonist CL 316,243 which enhances lipid beta-oxidation in addition to pyruvate-driven respiratory capacity (Figure 3D). Curiously, the enhanced lipid beta oxidation occurred independently of changes cellular glycolytic capacity or changes in key lipid metabolism proteins or lipid signaling capacity between brown adipocytes that overexpressed Nipsnap1 compared to GFP controls (Figure 3E-3F). Taken together, these results indicated that overexpression of Nipsnap1 in primary brown adipocytes significantly increases beta-oxidation capacity which is likely the driver for the increase in whole body energy expenditure and reduced weight gain in mice.

**Figure 3.**
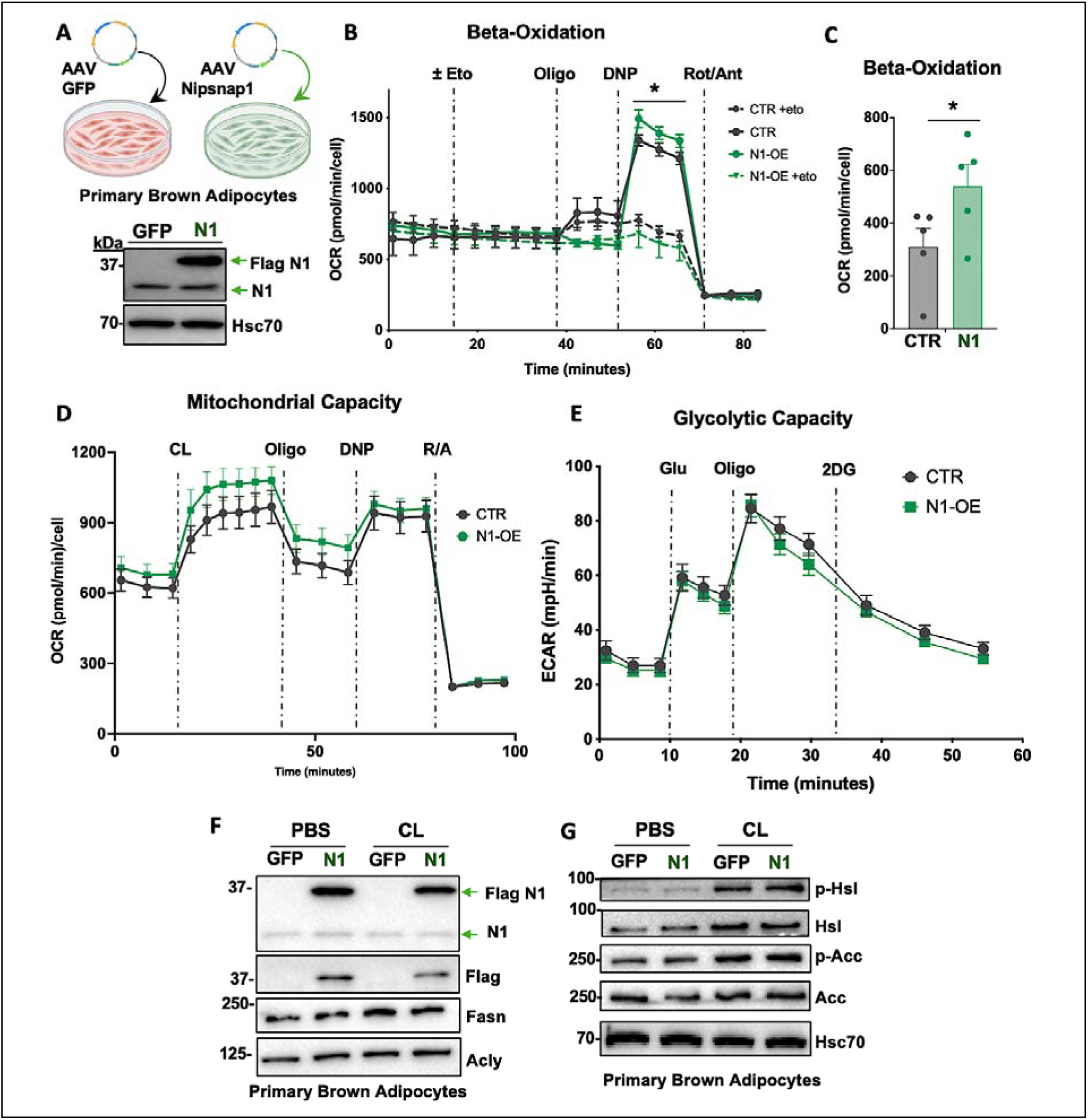
Overexpression of Nipsnap1 increases mitochondrial beta-oxidation capacity in brown adipocytes. **(A)** Upper panel: Schematic of AAV-GFP and AAV-Flag-N1 transduction into primary brown adipocytes. Lower panel: Representative Western blot probed for endogenous N1 and exogenous AAV- Flag-N1 in primary brown adipocytes. **(B)** Beta-oxidation-driven respiration measured by Seahorse Bioanalyzer in primary brown adipocytes transduced with AAV-GFP or AAV-Flag-N1 (n = 5). **(C)** Quantification of beta-oxidation basal respiration in primary brown adipocytes transduced with AAV-GFP or AAV-Flag-N1 (n = 5). **(D)** Seahorse oxygen consumption rates of primary brown adipocytes transduced with AAV-GFP or AAV-Flag-N1 (n = 8). **(E)** Extracellular acidification rates (ECAR) of primary brown adipocytes transduced with AAV-GFP or AAV-Flag-N1 (n = 10). **(F-G)** Representative Western blot of primary brown adipocytes transduced with AAV-GFP or AAV-Flag-N1 and treated with PBS or 10 µM CL for 20 minutes. Hsc70 serves as the loading control. Data are represented as mean ± SEM unless indicated otherwise. Significance is *p < 0.05 by Student’s t-test with multiple corrections by Bonferroni.

### Nipsnap1 is an integral component of the mitochondrial and peroxisomal beta-oxidation protein network

In our previous study, we demonstrated that Nipsnap1 is critical for functional lipid metabolism^8^. The protein interaction network of Nipsnap1 in brown adipocytes however is completely unknown. To map this network, we interrogated the protein binding partners of Flag-tagged Nipsnap1 compared to GFP treated controls in primary brown adipocytes by co- immunoprecipitation (Figure 4A and B). Immunoprecipitation of Nipsnap1 revealed interactions with several protein binding partners of which the most notable were interactions with lipid metabolism proteins in both the mitochondria and within peroxisomes (Figure 4C and D). We also pulled down the established Nipsnap1 interacting protein, Nipsnap2 confirming the reported localization of Nipsnap1 and its ability to translocate between the inner and outer mitochondrial membranes as a sensor of mitochondrial health^9^. We further show that Nipsnap1 binds to key mitochondrial lipid metabolism proteins such as the mitochondrial acylcarnitine carrier protein (slc25a20) suggesting that Nipsnap1 plays a role in the regulation of lipid import into the mitochondria (Figure 4E). We also show that Nipsnap1 binds to enoyl-CoA hydratase and 3- hydroxyacyl Co-A dehydrogenase (Ehhadh) which is one of the key enzymes in the peroxisomal beta-oxidation pathway (Figure 4E). These binding partners complement the emerging physiological role of Nipsnap1 where we demonstrate both through ablation and overexpression the importance of this regulatory factor in mitochondrial lipid metabolism in BAT.

**Figure 4.**
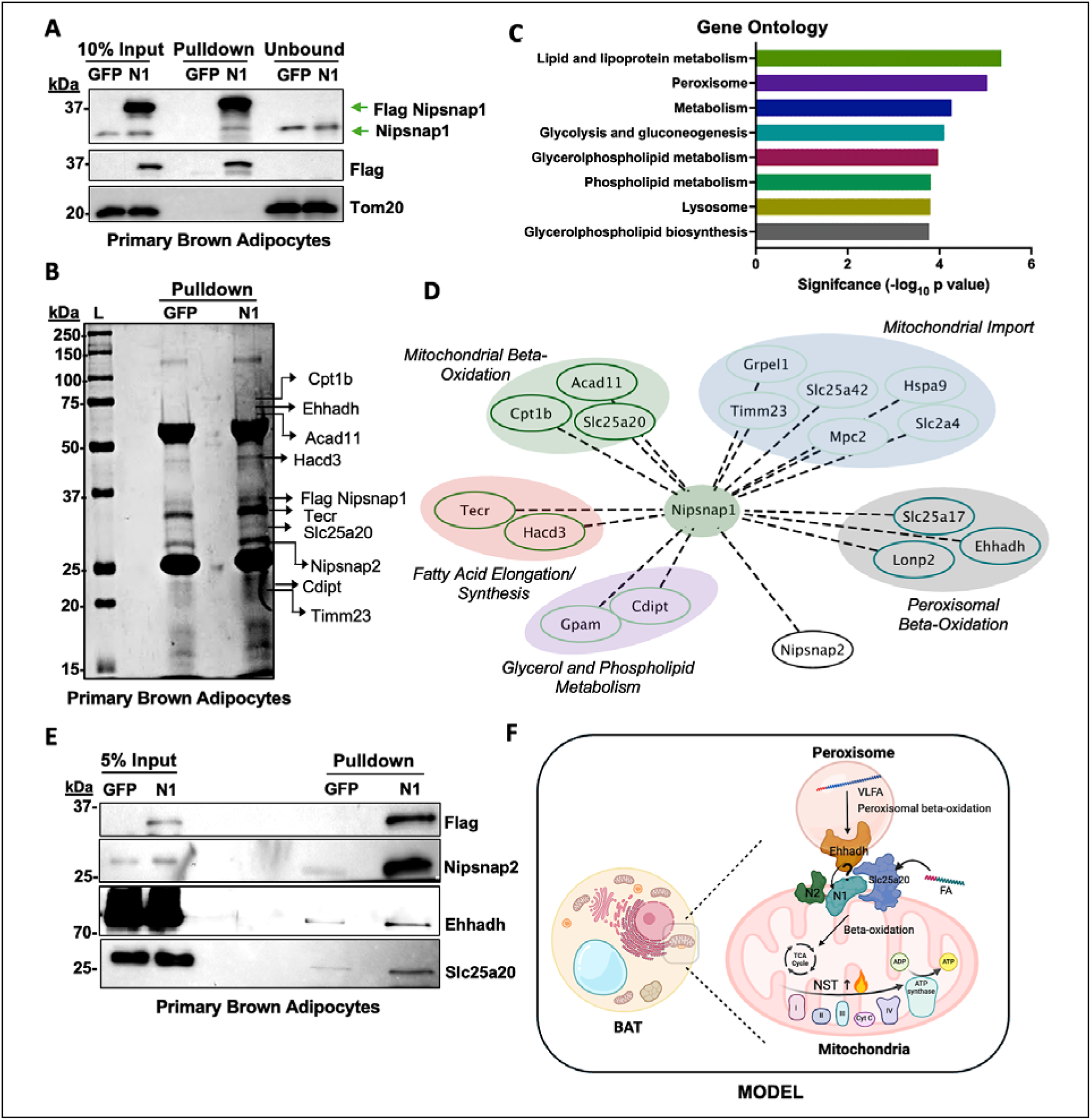
The protein interacting network of Nipsnap1 to mitochondrial and peroxisomal mediated beta-oxidation. **(A)** Representative western blot of Flag-tagged N1 immunoprecipitation from crude mitochondrial fractions of primary brown adipocytes transduced with AAV-GFP or AAV-Flag-N1. Tom20 serves as a loading control. **(B)** Coomassie blue-stained SDS-PAGE gel of anti-Flag immunoprecipitation products from primary brown adipocytes. L= ladder. **(C)** Gene ontology analysis of proteomic data from Flag-N1 pulldown samples. **(D)** Protein-protein interaction network of N1 in primary brown adipocytes, visualized using Cytoscape. **(E)** Representative immunoblot of N1 interacting proteins. **(F)** Proposed model of N1 function in brown adipocyte metabolism. N1 serves as a tether to interact with Ehhadh to enhance peroxisomal beta-oxidation of very long-chain fatty acids (VLFA), while also associating with Cpt1b on the mitochondrial membrane to facilitate medium- and short-chain fatty acid (FA) import and oxidation. These interactions promote mitochondrial beta-oxidation in brown adipocytes, ultimately increasing energy production through non-shivering thermogenesis (NST).

## DISCUSSION

We previously defined the mitochondrial protein Nipsnap1 as a critical regulator for the long- term maintenance of non-shivering thermogenesis in brown adipose tissue^8^. The ablation of Nipsnap1 in mice resulted in an impaired ability to maintain body temperature during cold exposure due to significant reductions in total energy expenditure. Mechanistically, we demonstrated that the loss of Nipsnap1 caused significant disruptions in lipid metabolism as evidenced by decreased expression of key lipid proteins and decreased mitochondrial beta- oxidation^8^. It was clear that Nipsnap1 was critical for lipid metabolism in BAT, but how Nipsnap1 integrated with this process to support the long-term activation of brown adipose tissue was unknown.

Previous research has demonstrated that Nipsnap1, in conjunction with Nipsnap2, localizes to both the inner and outer mitochondrial membrane, where it plays a critical role in recruiting autophagy factors. This recruitment is essential for the initiation and regulation of mitophagy, an autophagic process that selectively degrades damaged mitochondria^9^. By enhancing mitophagy, Nipsnap1 helps preserve mitochondrial integrity, thereby mitigating oxidative stress and maintaining overall mitochondrial health^9^. Previous research has also demonstrated that Nipsnap1 plays a role in mediating lipid metabolism in the liver by interacting with proteins involved in regulating long chain fatty acid and glycerol metabolism^10^, however, the mechanism and the binding protein network of Nipsnap1 in regulating lipid metabolism was unknown.

In this study, we set out to elucidate Nipsnap1’s role in lipid metabolism and energy homeostasis in brown fat. We generated an adipose-specific AAV construct to overexpress Nipsnap1 specifically in thermogenic adipocytes of mice. In alignment with the emerging role of Nipsnap1 as a critical regulator in lipid metabolism, we show that overexpression of Nipsnap1 in mice resulted in a metabolic shift towards lipid substrate utilization as evidenced by reduced respiratory exchange ratio contributing to the increase in total energy expenditure compared to control mice. This increase in energy expenditure correlated with enhanced protection of their body temperature during cold exposure and decreased body weight gain. The increased lipid metabolism exhibited by the Nipsnap1-OE mice is likely driven by brown adipocytes where we observed increased levels of beta-oxidation capacity in the primary cultured brown adipocytes that overexpressed Nipsnap1.

In this study, we mapped the first Nipsnap1 protein-protein interacting network. Nipsnap1 possesses an N-terminal mitochondrial targeting sequence, and we have demonstrated previously that this protein localizes in the mitochondria in brown adipocytes^11,12^. Therefore, it came as no surprise when Nipsnap1 exhibited strong binding affinities for mitochondrial proteins involved in beta-oxidation. The most robust binding partner of Nipsnap1 was solute carrier family 25 member 20 (Slc25a20), an acyl-carnitine transporter in the mitochondrial inner membrane that facilitates the transfer of medium-chain acyl-CoAs into the mitochondrial matrix for beta- oxidation^13^. An additional Nipsnap1 binding partner is carnitine palmitoyl transferase 1b (Cpt1b), the major isoform of Cpt1 family. Cpt1b is localized on the outer membrane of the mitochondria and is expressed in cardiomyocytes, muscle, and brown fat, and has been indicated as a rate-controlling enzyme of the long-chain fatty acid beta-oxidation pathway in the mitochondria^14,15^. Beta-oxidation takes place in both the mitochondria and the peroxisomes. In peroxisomes, very long chain fatty acids (VLFA) are broken down into long- and medium-chain fatty acids by peroxisomal beta-oxidation enzymes^16^. These processed long- and medium- chain fatty acids are then transported into the mitochondria, where fatty acids are further catabolized into acetyl-CoA by mitochondrial beta-oxidation enzymes^17^. Intriguingly, immunoprecipitation of Nipsnap1 also revealed strong association with several peroxisomal-mediated beta-oxidation enzymes, such as enoyl-coA hydratase and 3-hydroxyacyl coA dehydrogenase (Ehhadh). Both the hydratase and dehydrogenase activity of the Ehhadh protein catalyzes the third and fourth steps of peroxisomal beta oxidation respectively^18^. Nipsnap1 also bound to the Slc25a17 protein which functions as a peroxisomal transporter for multiple cofactors including CoA, FAD, FMN, and AMP which are essential cofactors to maintain redox homeostasis for beta-oxidation in the perioxisome^19–21^.

Given the fact that Nipsnap1 binds to both mitochondrial and peroxisomal lipid beta-oxidation proteins, it is possible that Nipsnap1 serves as a putative tether or mediator between mitochondria and peroxisomes. Given the reported localization of Nipsnap1 to the outer mitochondrial membrane and the fact that it can transition between the inner and outer mitochondrial membrane^9^, Nipsnap1 would be well-positioned to mediate an extracellular mitochondrial–peroxisomal connection. This is supported by our finding that Nipsnap1 binds to the acyl-CoA dehydrogenase 11 (Acad11) protein that exhibits a complex subcellular distribution in both mitochondria and peroxisomes, suggesting a role in coordinating very long-chain fatty acid oxidation between these organelles. The potential interaction of Acad11 with Nipsnap1 may facilitate this inter-organelle lipid crosstalk, enhancing efficiency to transport lipids between the mitochondria and the peroxisome^22,23^. Previous studies have shown that Nipsnap1 colocalizes with its homolog, Nipsnap2, on the mitochondrial outer membrane, playing a role in regulating mitochondrial homeostasis and the immune response^24^. This present study also identified a strong binding affinity between Nipsnap1 and Nipsnap2, supporting the hypothesis that Nipsnap1 may form a complex with Nipsnap2 on the mitochondrial membrane and function as a transporter to facilitate the import of lipids by looping very long-chain fatty acid metabolites processed by the peroxisomes and transferring them into the mitochondria as substrates for beta-oxidation (Figure 4F). Furthermore, there is increasing evidence supporting the notion that peroxisomes and mitochondria engage in vital, significant crosstalk, which is crucial for preventing peroxisomal dysregulation and accumulation of reactive oxygen species (ROS) stress^25,26^. Membrane contact sites (MCS) have emerged as a leading explanation for this constant material exchange between the organelles^27^. An MCS involves a bridge-like formation spanning the short distance between closely positioned mitochondria and peroxisomes, facilitating the direct transport of beta-oxidation substrates, ROS, and other materials. Multiple theories have been proposed to describe the composition of these MCS bridge formations. In *Saccharomyces cerevisiae*, proteins such as Pex11 and Mdm34 (part of the ER-mitochondrial tether complex) have been implicated^28^. In mammals, the ATP-binding cassette transporter 1 (Abcd1) has been identified as a potential component of these contact sites^29,30^. However, further research is required to determine whether this process is directly linked to Nipsnap1.

## Conclusions

We have identified that the Nipsnap1 protein interacts with key regulators of both peroxisomal and mitochondrial beta-oxidation pathways in BAT. This interaction enhances the utilization of lipids as energy substrates, thereby sustaining the long-term activation of brown fat and enhanced metabolic health in mice. Taken together, these results solidified our claim that Nipsnap1 is critical for lipid metabolism in BAT and could be a therapeutic target to protect against obesity-associated metabolic dysfunction.

## Methods

### Animals

All experimental procedures involving mice were conducted with approval from the Institutional Animal Care and Use Committee at Cornell University. Three-week-old wild-type male C57BL/6J mice (Jackson Laboratory, #000664) were maintained at room temperature (25 °C) with a 12-hour light/dark cycle and had unlimited access to food and water. For adeno-associated virus (AAV) transduction, five-week-old mice were anesthetized using isoflurane and tail-vein injected with virus overexpressing either eGFP or Flag-Nipsnap1 (N1) (Penn Vector Core) at a dose of 2.5x10^12^ viral genomes (vg) per mouse. The mice were then kept at room temperature for 14 days to allow the virus to infect the adipocytes before conducting further experiments. At the end of each experiment, the mice were euthanized with carbon dioxide. Following euthanasia, various tissues, including brown adipose tissue, inguinal adipose tissue, white adipose tissue, liver, and brain, were collected. Each tissue sample was promptly subjected to flash-freezing utilizing liquid nitrogen. The samples were then stored at a temperature of −80 °C for subsequent protein or RNA extraction.

### *In vivo* Indirect Calorimetry

Six-week-old C57BL/6J mice (Jackson Laboratory, #000664) infected with AAV GFP or AAV Nipsnap1 were individually housed in Promethion metabolic cages (Sable Systems International) on a 10/14-hour light/dark cycle with free *ad libitum* access to water and food. Mice were then acclimated in the metabolic cages at 25°C for 24 hours before the temperature was gradually ramped down to 6°C for the cold exposure experiment. Real-time metabolic parameters, such as energy expenditure (kcal/hr), oxygen consumption (VO2), carbon dioxide production (VCO2), locomotor activity (assessed by X-Y infrared beam breaks), and the respiratory exchange ratio (RER), were recorded at 3-minute intervals using the Sable System data acquisition software (IM-3 v.20.0.3). The collected raw data were processed with Sable System Macro Interpreter software and One-Click Macro Systems (v2.37), followed by further analysis using CalR software (v1.3).

### Cell Culture

Primary brown adipose tissue was harvested from three-week-old male wild-type C57BL/6J mice, then thoroughly minced with scissors for 5 minutes and digested with 15 mL of lysis buffer (123 mM NaCl, 5 mM KCl, 1.3 mM CaCl2, 5.0 mM Glucose, 100 mM HEPES, 4% BSA, and 1.5 mg/mL collagenase B) by incubating in a 37 °C constant shaking water bath for 30 minutes. After the digestion buffer was removed, the stromal vascular fraction (SVF) was resuspended in adipocyte culture media (DMEM/F12 with 10% FBS, 25 mM HEPES, and 1% PenStrep) and filtered through a 40 μm cell strainer, then centrifuged at 600 g for 5 minutes at 4 °C. The cells were subsequently resuspended in adipocyte culture media and plated on 10 cm polystyrene cell culture plates coated with 2% gelatin. After 48 hours, the cells were gently washed twice with PBS and replenished with fresh adipocyte culture media. All primary brown adipocytes were cultured at 37 °C with 5% CO2. For adipocyte differentiation, cells were seeded on polystyrene cell culture plates with 2% gelatin coating. The following day, 100% confluent cells were differentiated with DMEM/F12 (supplemented with 5 μg/mL insulin, 1 μM Rosiglitazone, 1 μM Dexamethasone, 0.5 mM Isobutylmethylxanthine, and 1 nM T3). After 48 hours, the medium was replaced with maintenance media (5 μg/mL insulin, 1 μM Rosiglitazone, and 1 nM T3) and replenished every two days. To achieve overexpression of either AAV-eGFP or N1, primary brown adipocytes were transduced with AAV at a multiplicity of infection (MOI) of 7.5x10^4^.

This transduction was performed over a 48-hour period, starting on Day 3 and ending on Day 5 of the differentiation process. By Day 7, the cells were fully differentiated and prepared for various experimental assessments. For acute CL 316,243 treatment, fully differentiated primary brown adipocytes were treated with 1 μM CL for 20 minutes.

### Cellular Respiration Assay

To assess cellular respiration in primary brown adipocytes, 20,000 cells were cultured and differentiated on XFe24 Seahorse cell culture plates coated with 2% gelatin, following a previously established protocol^8^. To measure oxygen consumption rates (OCR), fully differentiated cells were washed and incubated for 30 minutes in an unbuffered DMEM solution (4.5 g/L glucose, 4 mM glutamine, 100 mM pyruvate, and 2% fatty acid-free BSA, with the pH adjusted to 7.4 using NaOH). For measurements of acute CL-driven mitochondrial capacity, specific compound concentrations were used.: Port A: CL 316,243 (10 µM), and Port B: Oligomycin (4 µM), Port C: DNP (0.6 mM), Port D: Rotenone/Antimycin A (4 µM each).

To assess cellular beta-oxidation, fully differentiated cells were fasted overnight in non-glucose DMEM supplemented with 0.5 mM glucose, 1 mM glutamine, 0.5 mM L-carnitine, and 1% FBS. On the day of measurement, the cells were switched to unbuffered DMEM containing 2 mM glucose and 0.5 mM L-carnitine and incubated for 30 minutes. The compound concentrations were then as follows: Port A: Etomoxir (10 µM) or unbuffered DMEM, Port B: Oligomycin (4 µM), Port C: DNP (0.6 mM), Port D: Rotenone/Antimycin A (4 µM each).

For extracellular acidification rate (ECAR) measurements, fully differentiated cells were washed and incubated for 30 minutes in unbuffered DMEM supplemented with 2 mM L-glutamine. The cellular glycolysis stress test involved administering the following compounds: Port A: Glucose (25 mM), Port B: Oligomycin (4 µM), Port C: 2-Deoxy-D-glucose (2-DG) (50 mM). After each experiment, cells were stained with Hoechst, and cell numbers were quantified using the Lionheart FX automated microscope (Agilent) for normalization.

### Protein Extraction and Western Blotting

Tissue lysates were prepared using a 2% Sodium Dodecyl Sulfate (SDS) lysis buffer solution, supplemented with a complete protease inhibitor (ThermoFisher #A32963). The tissues were homogenized using the Qiagen TissueLyser II at 4 °C for 30 minutes. For cell lysates, adipocyte cultures were scraped into a 2% SDS lysis buffer with a protease inhibitor and incubated at 4 °C for one hour. The cells then underwent ultrasonic treatment using a Bioruptor (American Laboratory Trading) for 10 minutes, with 30-second intervals of sonication and rest. The lysates were centrifuged at maximum speed for 20 minutes to collect the protein supernatant. Protein concentrations in both tissue and cell lysates were determined using the BCA Protein Assay Kit (ThermoFisher #23227). The protein samples were mixed with 4× Laemmli buffer (Bio-Rad #161074) and heated at 37 °C for 5 minutes. These samples were then separated using SDS-polyacrylamide gel electrophoresis and transferred onto PVDF membranes. The membranes were blocked with 5% milk for one hour, washed with TBST, and incubated overnight with primary antibodies. Membranes were washed with TBST, incubated with secondary goat anti-mouse or anti-rabbit antibodies, and washed again with TBST before imaging with the FluorChem system (bio-techne).

### RNA Extraction and Real-Time Quantitative PCR

Adipocyte cells and tissue samples were lysed using Trizol reagent (Invitrogen) and the tissue samples underwent further homogenization utilizing the Qiagen TissueLyser II. Total RNA was extracted following the manufacturer’s protocol, and RNA concentration was measured using a Nanodrop spectrophotometer (ThermoFisher). Reverse transcription was performed with the High-Capacity cDNA Reverse Transcription Kit (ThermoFisher #4368813). Quantitative gene expression analysis was then conducted using the CFX384 Real-Time PCR System with SYBR Green (Bio-Rad #1725275) for real-time PCR.

### Subcellular Fractionation

Brown adipocyte tissues were isolated from mice and subcellular fractionation was performed by following the Qproteome Mitochondria Isolation Kit instructions (Qiagen #37612).

### Flag Tag Pull-Down Assay

Crude mitochondria were isolated from fully differentiated primary brown adipocytes as previously described^31^. The extracted mitochondrial pellets were lysed in RIPA buffer (50 mM Tris-HCl, pH 8.0, 150 mM NaCl, 0.1% SDS, 1% NP-40, 5 mM EDTA, and 0.5% sodium deoxycholate), and protein concentration was determined using the BCA Protein Assay Kit (ThermoFisher #23227).

Lysates containing eGFP and N1-Flag proteins were incubated overnight at 4 °C with Anti-FLAG® M2 Magnetic Beads (Millipore Sigma #M8823). The following day, the lysates were removed, and the magnetic beads were washed three times with TBS and rinsed three times with molecular-grade water. For comprehensive identification of binding partners, the magnetic beads underwent trypsin-on-beads digestion followed by mass spectrometry analysis at the Weill Cornell Proteomics and Metabolomics Core Facility. For further validation, the pulled-down magnetic beads were boiled in a 2× SDS buffer at 95 °C for ten minutes, and the protein lysates were resolved by SDS-PAGE, followed by Coomassie blue staining and western blotting.

### Gene Otology Enrichment and Protein Network Analysis

Gene Ontology (GO) enrichment analysis was conducted using the Enrichr database^32–34^ based on the binding proteins identified from the Flag-tagged pull-down assay, applying a cut-off threshold of fold changes greater than 2. The resulting protein-protein interaction network associated with the N1 protein was visualized using Cytoscape, incorporating only those binding proteins with a peptide abundance of more than three peptides.

### Antibody and primers

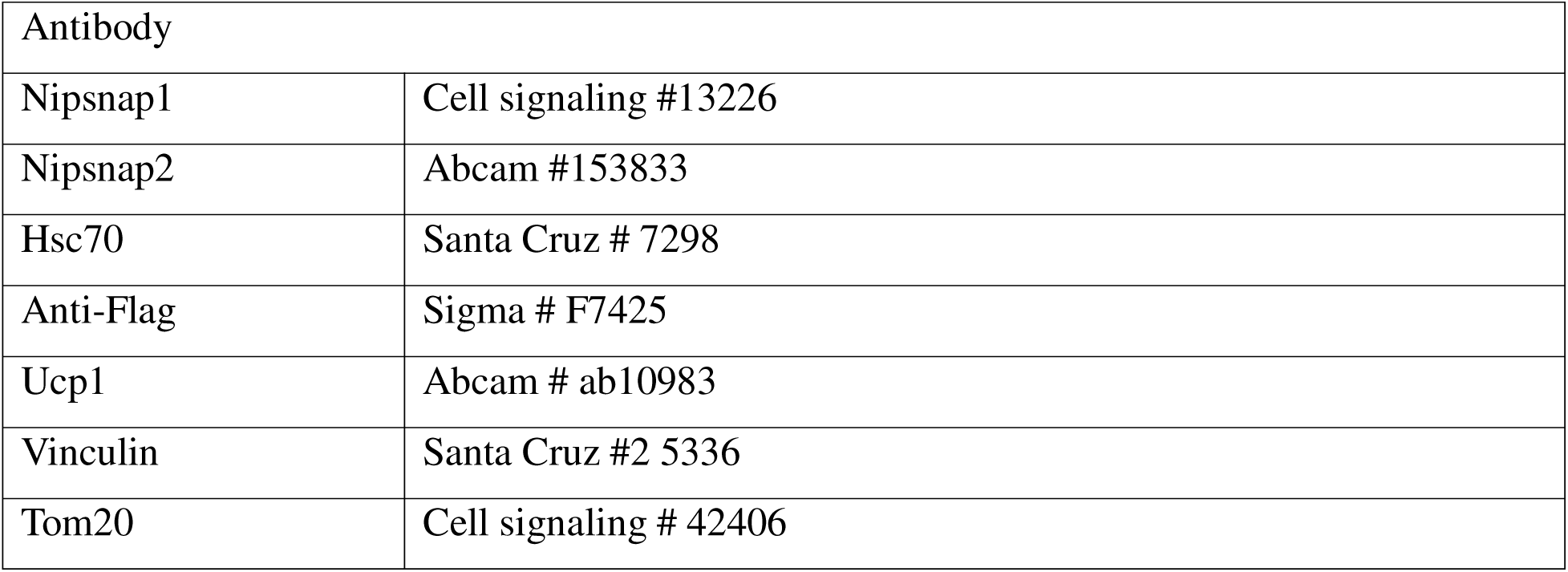
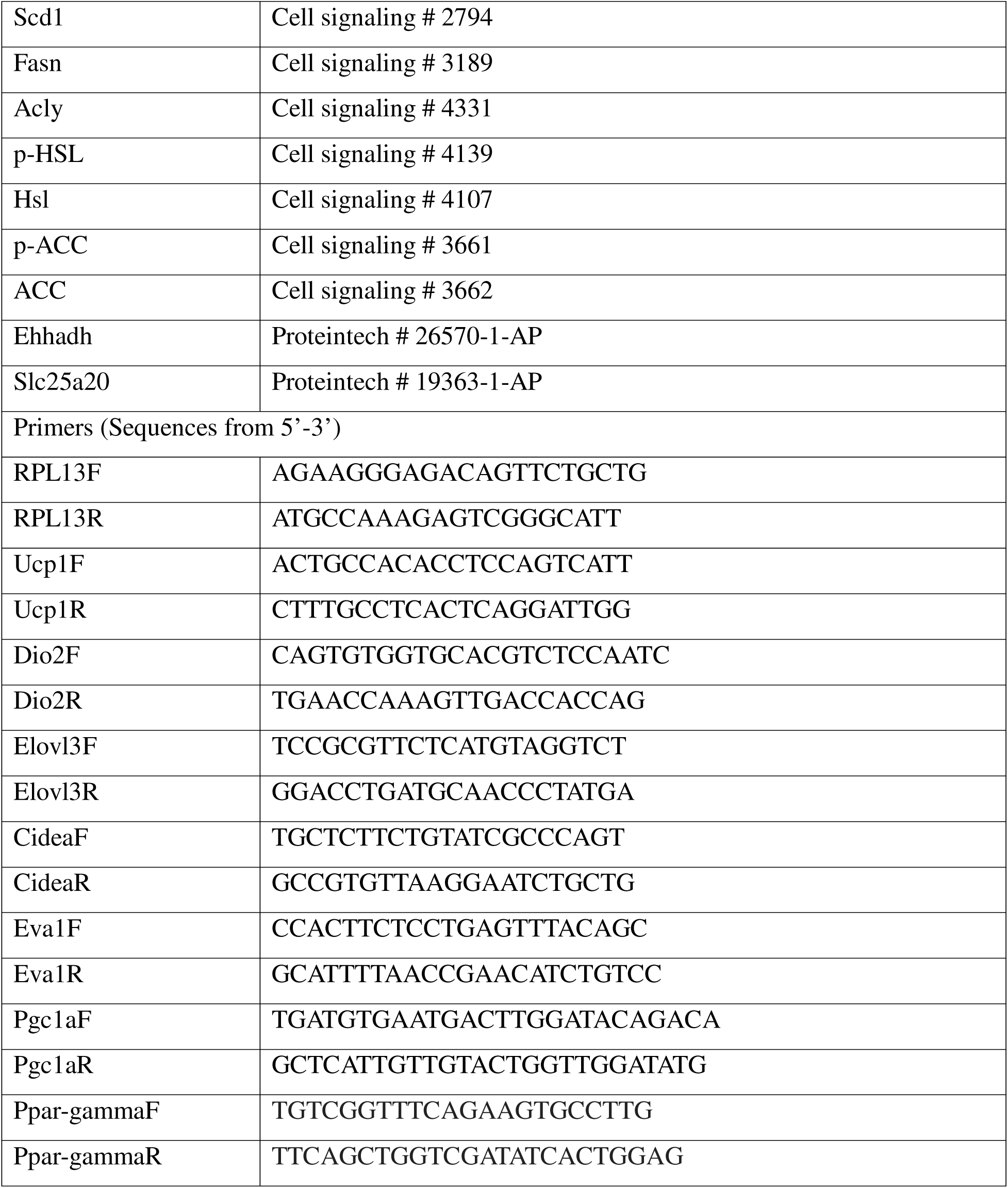

### Statistical analyses

Data are expressed as mean ± standard error of the mean (S.E.M.) unless otherwise noted. Statistical analyses were conducted using GraphPad Prism version 10. Group comparisons between two independent datasets were performed using unpaired two-tailed Student’s t-test. Metabolic measurements from Promethion were analyzed via analysis of covariance (ANCOVA) using CalR version 1.3, following established protocols ^35,36^

## Acknowledgements

Illustrations were created using BioRender. We thank the Penn Vector Core facility for the generation of the Nipsnap1 and control AAV8 viral particles. We also sincerely thank the Progressive Assessment of Therapeutics (PATh) at Cornell University for the tail vein injections of the AAV viral particles.

## Ethics declarations

Not applicable.

## Consent for publication

Not applicable.

## Author information

Authors and Affiliations

Pei-Yin Tsai, Yue Qu, Claire Walter, Yang Liu and Joeva Barrow-Division of Nutritional Sciences, Cornell University, Ithaca NY 14850, USA

Chloe Cheng- Department of Veterinary Medicine, Cornell University, Ithaca NY 14850, USA

## Author Contributions

The project was conceptualized by J.J.B. and P.-Y.T. and Y.L.; methodology: P.-Y.T., and Y.Q and Y.L.; investigation: P.-Y.T., Y.Q., Y.L., C.W., C.C., and J.J.B.; global analyses: P.-Y.T., and J.J.B.; writing: J.J.B. and P.-Y.T.; supervision: J.J.B. All authors have read and agreed to the published version of the manuscript.

## Corresponding author

Correspondence to Joeva Barrow, Ph.D., R.D. 244 Garden Avenue Savage Hall Room 122, Cornell University, Ithaca, NY 14850, USA. Phone: +1 607 955 9425.

## Data Availability Statement

Data described in the manuscript will be made available upon request.

